# Comparative studies of the seven human coronavirus envelope proteins using topology prediction and molecular modelling to understand their pathogenicity

**DOI:** 10.1101/2021.03.08.434384

**Authors:** Dewald Schoeman, Ruben Cloete, Burtram C. Fielding

## Abstract

Human (h) coronaviruses (CoVs) 229E, NL63, OC43, and HKU1 are less virulent and cause mild, self-limiting respiratory tract infections, while SARS-CoV, MERS-CoV, and SARS-CoV-2, are more virulent and have caused severe outbreaks. The CoV envelope (E) protein, an important contributor to the pathogenesis of severe hCoVs infections, may provide insight into this disparate severity of the disease. Topology prediction programs and 3D modelling software was used to predict and visualize structural aspects of the hCoV E protein related to its functions. All seven hCoV E proteins largely adopted different topologies, with some distinction between the more virulent and less virulent ones. The 3D models refined this distinction, showing the PDZ-binding motif (PBM) of SARS-CoV, MERS-CoV, and SARS-CoV-2 to be more flexible than the PBM of hCoVs 229E, NL63, OC43, and HKU1. We speculate that the increased flexibility of the PBM may provide the more virulent hCoVs with a greater degree of freedom, which can allow them to bind to different host proteins and can contribute to a more severe form of the disease. This is the first paper to predict the topologies and model 3D structures of all seven hCoVs E proteins, providing novel insights for possible drug and/or vaccine development.

## INTRODUCTION

Of the seven human (h) coronaviruses (CoVs) identified, the recent three, severe acute respiratory syndrome (SARS)-CoV, Middle East respiratory syndrome (MERS)-CoV, and SARS-CoV-2, are the most virulent and have caused severe outbreaks in the last two decades (Broadbent, 2020; Hewings-Martin, 2020). The other four hCoVs, hCoV-229E, −NL63, −OC43, and −HKU1, are less virulent and circulate continuously within the human population, peaking seasonally in different countries all over the world (Aldridge, et al., 2020; Cui, et al., 2011; Edridge, et al., 2020; Gaunt, et al., 2010; Killerby, et al., 2018; Lau, et al., 2006; Su, et al., 2016; Zeng, et al., 2018). They generally cause less severe acute respiratory tract infections, such as the common cold, in immunocompetent persons, but can become more severe in immunocompromised persons, the elderly, and those with chronic, underlying medical conditions (Killerby, et al., 2018; Liu, et al., 2020; Trombetta, et al., 2016). It is unclear, though, why these four hCoVs generally cause less severe disease in immunocompetent persons than the three more virulent hCoVs.

The disease severity and immunopathology of SARS-CoV and SARS-CoV-2 infections have largely been attributed to the envelope (E) protein (DeDiego, et al., 2007; Fett, et al., 2013; Jimenez-Guardeño, et al., 2014; Lamirande, et al., 2008; Nieto-Torres, et al., 2014; Nieto-Torres, et al., 2015; Regla-Nava, et al., 2015; Schoeman and Fielding, 2020; Xia, et al., 2020). No research, however, has demonstrated whether this is also the case for MERS-CoV infections. Despite the poor sequence identity of the E protein between the seven hCoVs, certain functional aspects, such as its postsynaptic density protein 95 (PSD95)/Drosophila disc large tumour suppressor (Dlg1)/zonula occludens-1 protein (zo-1) (PDZ)-binding motif (PBM) and ion-channel (IC) activity, appear to remain conserved and have both been implicated in the pathogenesis and clinical presentation of SARS-CoV and SARS-CoV-2 infections (Huang, et al., 2020; Nieto-Torres, et al., 2014; Nieto-Torres, et al., 2015; Schoeman and Fielding, 2020; Teoh, et al., 2010; Wang, et al., 2020; Wolff, et al., 2020; Xia, et al., 2020). The SARS-CoV E protein can interact with, among others, host cell proteins syntenin and the protein associated with *Caenorhabditis elegans* lin-7 protein 1 (PALS1) in the cytoplasm of host cells through its C-terminal PBM, highlighting the importance of its topology (Jimenez-Guardeño, et al., 2014; Teoh, et al., 2010). Recently, a peptide mimicking the PBM of SARS-CoV-2 E was also shown to be capable of interacting with PALS1 (Toto, et al., 2020). Despite some experimental evidence, the topology of the hCoV E protein remains largely under debate and data is limited to the prototypic SARS-CoV and to SARS-CoV-2 (Arbely, et al., 2004; Duart, et al., 2020; Khattari, et al., 2006; Nieto-Torres, et al., 2011; Yuan, et al., 2006). The topology of the E protein for the less virulent hCoVs, however, is unknown with no evidence to suggest whether or not it adopts a topology similar to the more virulent hCoVs. Similarly, the three-dimensional (3D) structure of the less virulent hCoVs has not been experimentally resolved to date. To date, the only hCoV E protein structure that has been resolved experimentally is that of SARS-CoV E for which two templates, viz. 5×29 and 2mm4 are available, both spanning residues 8-65 of the protein (Li, et al., 2014; Surya, et al., 2018). The SARS-CoV E protein also has other functions and is involved in other host cell process based on its interaction with different host cell proteins (Schoeman and Fielding, 2019). Given the various functions of E protein in the CoV life cycle, it is suggested to be capable of adopting multiple topologies, depending on the respective function (Kuo, et al., 2007).

The complexity and largely hydrophobic nature of membrane proteins, such as the CoV E protein, presents a challenge in studying their structure and dynamics experimentally (Arbely, et al., 2004; Latek, et al., 2019; Wu, et al., 2003). In the absence of experimental data, bioinformatics software tools may provide useful answers in dissecting the topologies, which can then be explored experimentally. Recently, Duart, et al. (2020) used the prediction programs ΔG predictor, TMHMM, MEMSAT-SVM, TMpred, HMMTop, Phobius, TOPCONS to predict the topology of SARS-CoV-2 E protein, and validated the topology experimentally. The N-terminus of the SARS-CoV-2 E protein was oriented to the lumen of the endoplasmic reticulum-Golgi intermediate compartment (ERGIC) while the C-terminus was located cytoplasmically – a topology that would enable the C-terminal PBM to interact with host cell proteins in the cytoplasm. Since there is very little consensus on the topology of the E protein of more virulent hCoVs and none for the less virulent hCoVs, this study aims to employ the prediction programs used by Duart, et al. (2020) to predict the topology of the E protein for all seven hCoVs. Defining the membrane topology of the less virulent hCoVs can allude to whether they can interact with host cell proteins in a way similar to what the more virulent hCoVs do. This may provide insight into understanding the mechanism behind why hCoVs-229E, −NL63, −OC43, and −HKU1 are less virulent than their more virulent and recently emerging counterparts. It can also provide more insight into whether all hCoVs share a similar mechanism of assembly and release (Ruch and Machamer, 2012). The 3D structural models will provide insight into the pathogenicity of the virulent hCoVs by predicting the secondary and tertiary structural fold of the E protein’s PBM and thereby, establishing a structure-function relationship between the pathogenicity of the more virulent hCoVs vs the less virulent forms of hCoVs.

## METHODS

### Topology prediction of hCoV E protein

The amino acid sequences for the E protein of all seven hCoVs were obtained from the UniProt KB database (https://www.uniprot.org/) and only complete and “Reviewed” sequences were selected. The topology of the E proteins was predicted using the prediction programs ΔG Predictor (Hessa, et al., 2007) (http://dgpred.cbr.su.se/), TMHMM (Sonnhammer, et al., 1998) (http://www.cbs.dtu.dk/services/TMHMM/), MEMSAT-SVM (Nugent and Jones, 2009) (http://bioinf.cs.ucl.ac.uk/psipred/), TMpred (Hofmann, 1993) (https://embnet.vital-it.ch/software/TMPRED_form.html), HMMTop (Tusnady and Simon, 1998; Tusnady and Simon, 2001) (http://www.enzim.hu/hmmtop/), Phobius (Käll, et al., 2004) (http://phobius.sbc.su.se/), and TOPCONS (Tsirigos, et al., 2015) (http://topcons.net/). For each program, the user-adjustable parameters were kept at their default settings.

### Molecular modelling

Three-dimensional protein structures were constructed for all seven hCoV E proteins using MODELLER software to compare the structural features between the more virulent and less virulent forms (Eswar, et al., 2008; Sali and Blundell, 1993).

### Template selection and model construction

Two nuclear magnetic resonance (NMR)-resolved structures for SARS-CoV E (PDBID: 2mm4 and PDBID: 5×29) were obtained from the protein data bank (PDB) (Li, et al., 2014; Surya, et al., 2018). These structures were used to generate full-length 3D models for SARS-CoV (template: 5×29), SARS-CoV-2 (template: 5×29), MERS-CoV (template: 2mm4), and hCoVs 229E (template: 5×29), NL63 (template: 229E protein model), OC43 (template: 5×29), and HKU1 (template: 229E protein model). The routinely used python script, align2d.py, in MODELLER was used to perform an alignment prediction between each of the hCoV E protein sequences and the respective template sequence (Eswar, et al., 2008; Sali and Blundell, 1993). Thereafter, a 3D model was built for each hCoV E protein using the model-ligand.py script (Eswar, et al., 2008; Sali and Blundell, 1993). As a validation step, we also used the protein structure prediction server, ITASSER, to generate 3D structures for the seven hCoV E proteins and compared the structures to the MODELLER predicted structures (Zhang, 2008).

### Quality assessment

The phi and psi dihedral angle parameters for the Ramachandran plot was calculated for each of the predicted protein models using PROCHECK webserver (Laskowski, et al., 1993). The root mean square deviation analysis was done by performing structural alignment between the predicted model structure and the homologous template protein. This was done to assess if any structural deviation exists within the main chain atoms of the two protein structures. The predicted structures were visualized using PyMol molecular graphics software (DeLano, 2002).

## RESULTS

### Predicted topology of hCoV E protein

The output from each prediction program for the E protein of each hCoV can be found in the supplementary information (Tables S1–S7). The predicted topology of the E protein for all seven hCoVs exhibits little consensus, with only a marginal similarity within the more virulent hCoVs and the less virulent hCoVs, respectively (Table 1). While some of the more virulent and the less virulent hCoV E proteins have been predominantly predicted to possess two transmembrane domains (TMDs), the majority of the E proteins were predicted to have only one TMD.

**Table 1.**
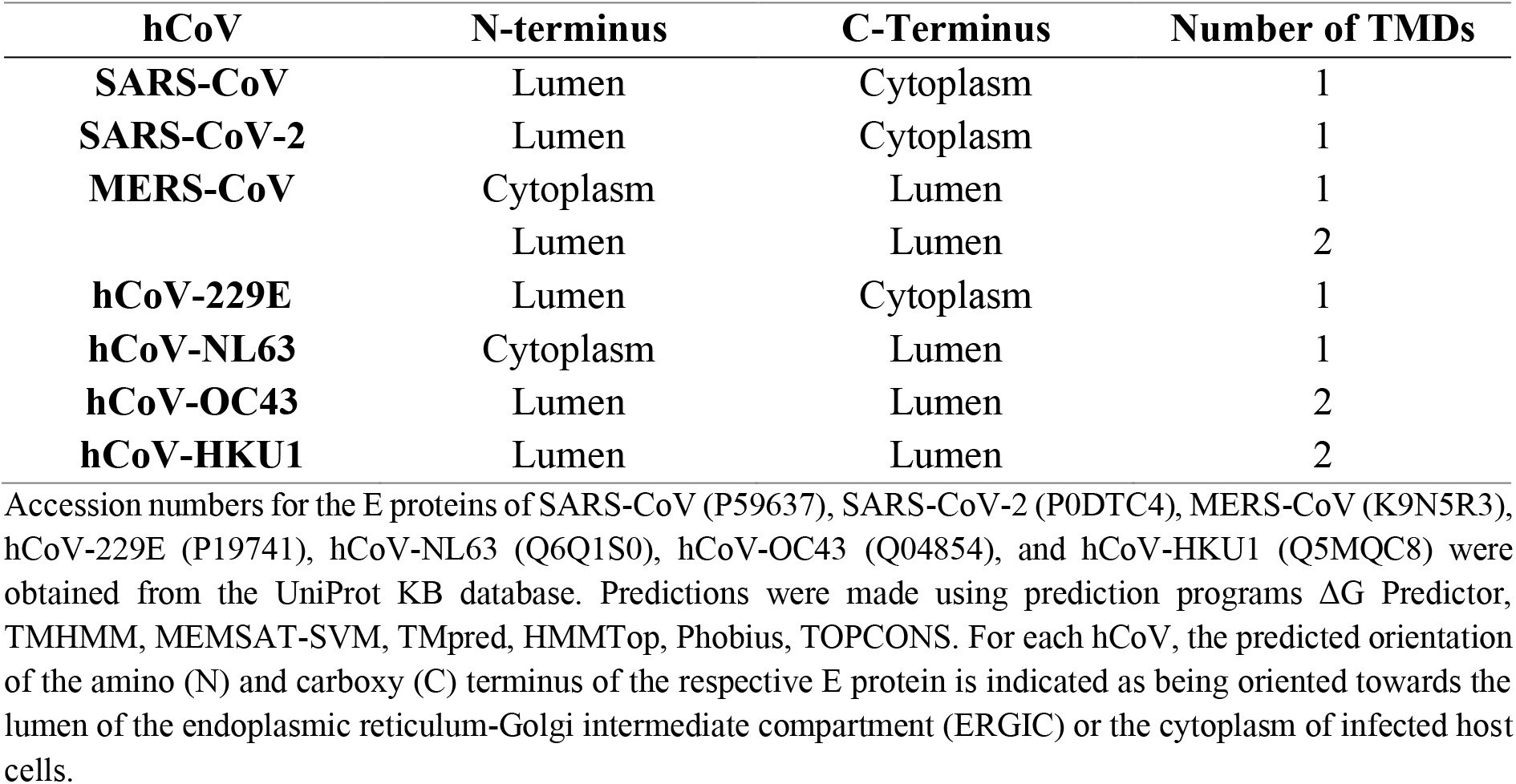
The predominant predicted topologies and number of transmembrane domains (TMDs) for all seven human (h) coronavirus (CoV) envelope (E) proteins.

#### Topology of virulent hCoV E proteins

Both SARS-CoV E and SARS-CoV-2 E were predicted to have one TMD with a lumenal N-terminus (N_lumen_) and cytoplasmic C-terminus (C_cyto_), while MERS-CoV E was predicted to adopt a contrasting topology of either a N_cyto_/C_lumen_ with a single TMD, or a N_lumen_/C_lumen_ with two TMDs, with an equal number of predictions for each topology (Table S3).

#### Topology of less virulent hCoV E proteins

The predicted topologies of the less virulent hCoVs were almost equally as inconsistent as the predicted topologies of the more virulent hCoVs. While hCoVs OC43 and HKU1 predominantly exhibited the same predicted topologies in having two TMDs and both N_lumen_/C_lumen_ orientations, hCoVs 229E and NL63 did not share this topology, nor were the respective topologies of the latter two similar. The hCoV-229E E protein was predominantly predicted to adopt a N_lumen_/C_cyto_ topology with a single TMD, while the hCoV-NL63 E protein was predicted to exhibit a N_cyto_/C_lumen_ topology also with a single TMD (Table 1).

### 3D models of hCoV E protein

#### Virulent hCoVs: SARS-CoV, SARS-CoV-2, and MERS-CoV

The SARS-CoV E protein shared 91% sequence identity with the amino acid sequence of template 5×29 (Figure S1A). The 3D model showed three α-helices and four coil regions, with the C-terminal PBM adopting a flexible coil region (Figure 1A). Quality assessment of the model revealed that 90% of the residues were in the most favoured regions of the Ramachandran plot and 1.4% were in the disallowed regions. The RMSD analysis showed a 0.698Å difference between SARS-CoV E and the template 5×29.

**Fig. 1.**
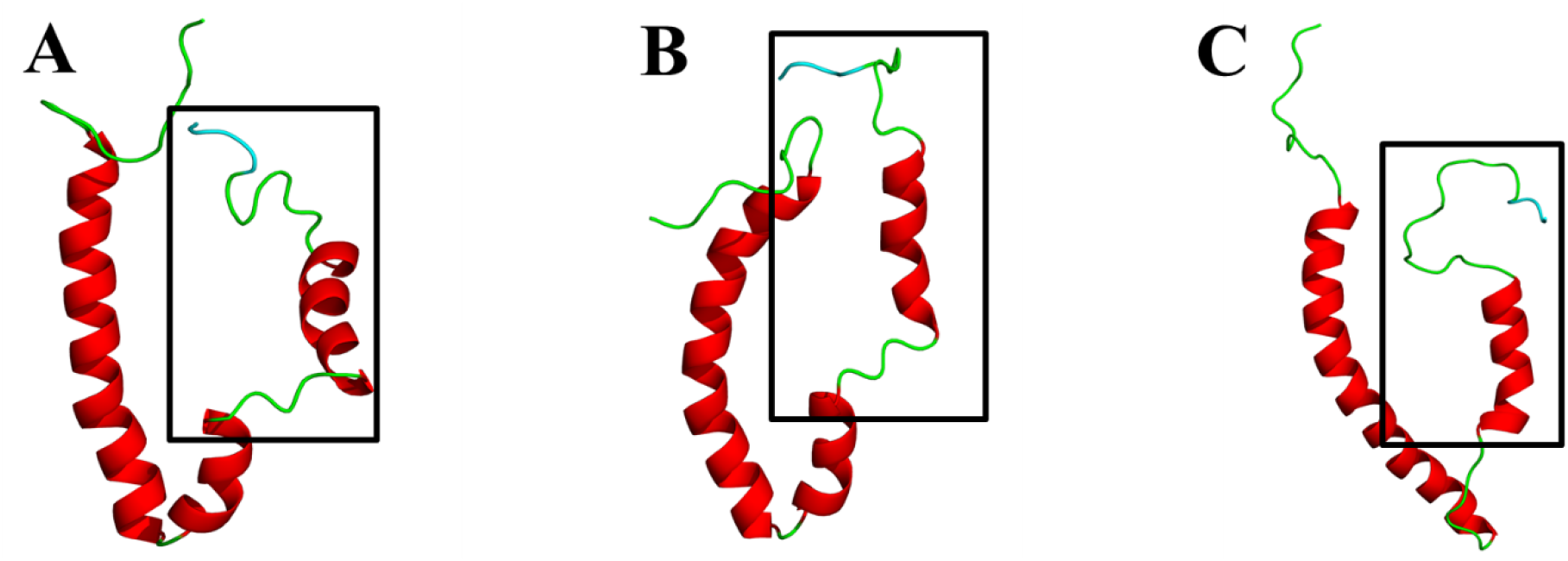
Cartoon representation of the three-dimensional (3D) models of the envelope (E) protein of the more virulent hCoVs. **A.** SARS-CoV E. **B.** SARS-CoV-2 E. **C.** MERS-CoV E. Models were generated using MODELLER software and based on two nuclear magnetic resonance (NMR)-resolved structures for SARS-CoV E (PDBID: 5×29 and PDBID: 2mm4) obtained from the protein data bank (PDB) (Eswar, et al., 2008; Li, et al., 2014; Sali and Blundell, 1993; Surya, et al., 2018). Models of the SARS-CoV and SARS-CoV-2 E proteins were generated from the 5×29 template, whereas the MERS-CoV E protein was generated from 2mm4 template. The PDZ-binding motif (PBM) is coloured in cyan and the squared region shows the variable C-terminal domain.

Similarly, SARS-CoV-2 E shared 91% sequence identity with template 5×29 (Figure S1B). The 3D model also showed three α-helices and four coil regions, with the PBM adopting a flexible coil region (Figure 1B). Quality assessment revealed that 88.6% of the residues were in the most favoured regions of the Ramachandran plot and 0% were in the disallowed regions. The RMSD analysis showed a 2.155Å difference between SARS-CoV-2 E and template 5×29.

The MERS-CoV E protein shared a 35% sequence identity to template 2mm4 (Figure S1C). While the 3D model showed only two α-helices and three coil regions, the PBM also adopted a flexible coil region (Figure 2C). Quality assessment indicated that 95.8% of the residues were in the most favoured regions of the Ramachandran plot and 1.4% were in the disallowed regions. The RMSD analysis between MERS-CoV E and the homologous template structure 2mm4 indicated a difference of 1.458Å.

**Fig. 2.**
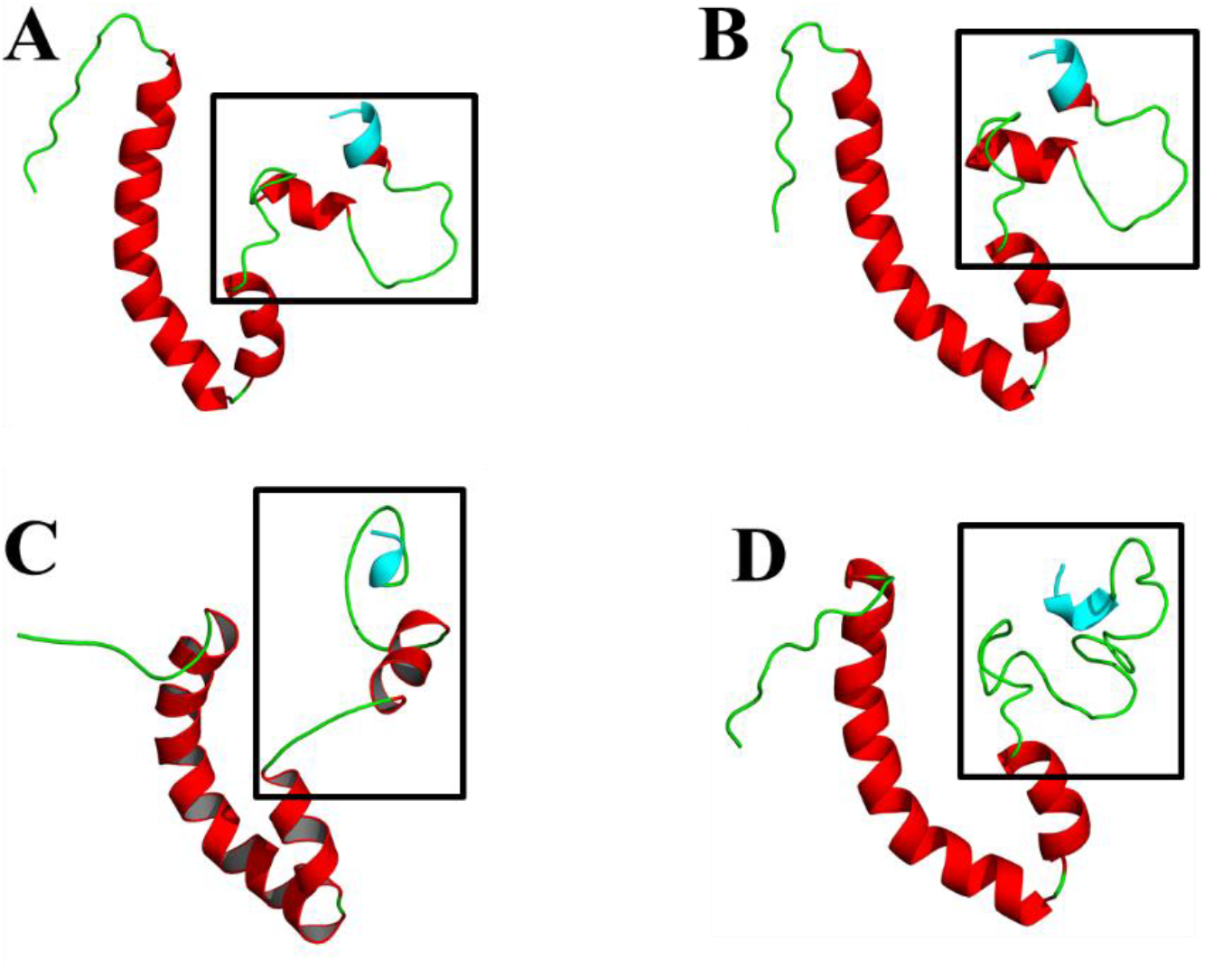
Cartoon representation of the three-dimensional (3D) models of the envelope (E) protein of the less virulent hCoVs. **A.** hCoV-229E E. **B.** hCoV-NL63 E. **C.** hCoV-OC43 E. **D.** hCoV-HKU1 E. Models were generated using MODELLER software and based on two nuclear magnetic resonance (NMR)-resolved structures for SARS-CoV E (PDBID: 2mm4 and PDBID: 5×29) obtained from the protein data bank (PDB) (Eswar, et al., 2008; Li, et al., 2014; Sali and Blundell, 1993; Surya, et al., 2018). Models of the hCoV-NL63 and −OC43 E proteins were generated from the 5×29 template, whereas models of the hCoV-229E and −HKU1 E proteins were generated from a 229E template. The PDZ-binding motif (PBM) is coloured in cyan and the squared region shows the variable C-terminal domain.

#### Non-virulent hCoVs: hCoV-229E, −NL63, −OC43, and −HKU1

The hCoV-229E E protein shared a 29% sequence identity with 5×29 amino acid sequence (Figure S2A). While the 3D model showed four α-helices and four coil regions, the C-terminal PBM adopted one α-helical turn conformation with reduced flexibility (Figure 2A) and Val74 shared conservation with Val78 from 5×29. Quality assessment of the model indicated that 91.5% of the residues were in most favoured regions of the Ramachandran plot and 0% were in the disallowed regions. The RMSD analysis between the hCoV-229E E protein and the homologous template structure 5×29 indicated a 1.776Å difference.

The hCoV-NL63 E protein shared 47% sequence identity with the amino acid sequence of the hCoV-229E E protein. (Figure S2B). Similar to E of hCoV-229E, the 3D model of hCoV-NL63 showed four α-helices and four coil regions, where the PBM adopted one α-helical turn conformation with reduced flexibility (Figure 2B), and conservation of Val74 was shared with Val74 from the E protein of hCoV-229E. Quality assessment indicated that 91% of the residues were in most favoured regions of the Ramachandran plot and 0% were in the disallowed regions. The RMSD analysis between hCoV-NL63 E and the homologous template structure (229E protein model) indicated a 1.217Å difference.

The hCoV-OC43 shared 22% sequence identity with the 5×29 amino acid sequence (Figure S2C). This structure also showed four α-helices and four coil regions, the PBM also adopted one α-helical turn conformation with reduced flexibility (Figure 2C), and Val81 shared conservation with Val78 from template 5×29. Quality assessment indicated that 90.8% of the residues were in the most favoured regions of the Ramachandran plot and 1.3% were in the disallowed regions. The RMSD analysis between hCoV-OC43 E and the homologous template structure 5×29 indicated a 3.328Å difference.

The hCoV-HKU1 E protein shared 27% sequence identity with the E protein amino acid sequence of hCoV-229E (Figure S2D). This model showed only three α-helices and three coil regions, while the PBM also adopted one α-helical turn conformation with reduced flexibility (Figure 2D), and Ile82 shared conservation with Ile75 from hCoV-229E E protein. Quality assessment indicated that 89.2% of the residues were in most favoured regions of the Ramachandran plot and 1.4% were in the disallowed regions. The RMSD analysis between hCoV-HKU1 E and the homologous template structure 229E indicated a 5.343Å difference.

## DISCUSSION

Despite the differences in the predicted number of TMDs for all seven hCoVs, the majority of hCoVs were predicted to have only one TMD, the length and number of which is largely consistent with reports for E in other CoVs (Lim and Liu, 2001; Liu, et al., 2007; Nieto-Torres, et al., 2011; Torres, et al., 2007; Torres, et al., 2005; Ye and Hogue, 2007; Yuan, et al., 2006). The predicted N_lumen_ and C_cyto_ topology of both the SARS-CoV E and SARS-CoV-2 E is consistent with what has been experimentally determined for the topology of untagged SARS-CoV E and c-myc-tagged SARS-CoV-2 E (Duart, et al., 2020; Nieto-Torres, et al., 2011). This similarity in topology is to be expected given the approximate 94% similarity in E protein amino acid sequence (Grifoni, et al., 2020; Schoeman and Fielding, 2020).

Given the lack of experimental evidence for the topology of MERS-CoV E, the reason(s) for and/or consequence(s) of these predicted topologies is not known. It may, however, provide some insight into why the human-human transmission of MERS-CoV is less effective than that of SARS-CoV and SARS-CoV-2 (Drosten, et al., 2014; Grijalva, et al., 2020; Wang, et al., 2020; Wilson-Clark, et al., 2006). The topology of membrane proteins, such as the E protein, plays an important part in their function, with certain functions, such as viral assembly and release of virions, being dependent on protein topology (Ruch and Machamer, 2012; Seppälä, et al., 2010; White, et al., 2015). The exact role of the CoV E protein in the assembly and release of newly formed virions was recently made clearer from the study of the infectious bronchitis virus (IBV) E protein (Westerbeck and Machamer, 2019). A low-molecular-weight (LMW), likely monomeric, form of IBV E increased the lumenal pH of the Golgi network employed by CoVs to assemble and egress from infected cells. This increased Golgi pH likely protects the spike (S) protein from premature cleavage, preventing the release of the trimer subunits prior to receptor binding. Interestingly, this disruption of the host secretory pathway was mediated by the E protein independent of its ion-channel activity, likely requiring interaction between the cytoplasmic tail of IBV E and a host protein capable of altering the pH of the Golgi network. Without the C-terminus of IBV E exposed to the cytoplasm, interaction with the necessary host protein would be unlikely and could result in the production of impaired, possibly non-infectious virions (Corse and Machamer, 2000; Westerbeck and Machamer, 2019). Neither of the predicted MERS-CoV E topologies reflects a cytoplasmic C-terminus, suggesting that MERS-CoV E might likely not be capable of interacting with a host protein in a way similar to IBV E to alter the host secretory pathway for the effective release of infectious virions. If MERS-CoV E does, in fact, adopt either of these topologies, it could explain why the virus is less effective at human-human transmission. Interestingly, an early X-ray scattering study suggested that E adopts a helical hairpin topology, whereas subsequent studies of solution NMR consistently demonstrated a single-span topology in which E was bound to several detergent micelles (Arbely, et al., 2004; Li, et al., 2014; Pervushin, et al., 2009; Surya, et al., 2018). This demonstrates the need to be cautious about which technique is used to determine E topology and take into account the environment to which E is exposed during the study. Nevertheless, the predicted topologies may merely reflect the ability of the E protein to adopt multiple topologies to perform specific functions (Kuo, et al., 2007; Westerbeck and Machamer, 2015; Westerbeck and Machamer, 2019). The lack of experimental evidence also impedes the reason(s) for and consequences of the predicted topologies.

The severe lack of experimental data on the E protein of the less virulent hCoVs makes it especially difficult to validate both the accuracy of the predicted topologies and their relevance. The requirement of the C-terminally located PBM of SARS-CoV E in the cytoplasm has been well-established as crucial to its pathogenesis, while no distinct function has been ascribed to the N-terminus (Castaño-Rodriguez, et al., 2018; Jimenez-Guardeño, et al., 2014; Teoh, et al., 2010). Thus far, only one study has investigated a possible function of the protein’s N-terminus whereby a chimeric SARS-CoV E protein was shown to contain possible Golgi-targeting information in its N-terminus (Cohen, et al., 2011). It should, however, be confirmed using the full-length protein as well.

A N_lumen_/C_cyto_ topology enables the SARS-CoV E PBM to interact with host cell proteins such as syntenin and PALS1. More recently, SARS-CoV-2 E has also been shown to interact with PALS1 in a similar manner, very likely since it adopts the same topology as SARS-CoV E (Duart, et al., 2020; Nieto-Torres, et al., 2011). If the C-terminus of SARS-CoV E was not oriented towards the cytoplasm, it would not be able to interact with these host proteins. Therefore, since the E proteins of the less virulent hCoVs NL63, OC43, and HKU1 were predicted to have a lumenal C-terminus, it might not allow for interaction with the same host proteins as SARS-CoV E; a feature that could likely explain why these hCoVs cause less severe infections compared to SARS-CoV. The same, however, cannot be said for the predicted topology of the less virulent hCoV-229E E protein. Despite sharing a topology similar to SARS-CoV E and SARS-CoV-2 E, hCoV-229E is self-limiting and causes less severe symptoms than the former two hCoVs (Lew, et al., 2003; Pappas, et al., 2008; Poutanen, 2018). Nevertheless, given the various functions of E in the CoV life cycle, it has also been proposed to assume multiple topologies, depending on the function (Nieto-Torres, et al., 2011; Ruch and Machamer, 2012; Yuan, et al., 2006). Since only a small proportion of the E protein produced during an infection is incorporated into CoV virions, it stands to reason that one topology might be adopted to facilitate the assembly of new viral progeny, while another is involved in the release through the secretory pathway (Westerbeck and Machamer, 2019). This was attempted where the IBV E protein was produced to adopt either a transmembrane topology, with one TMD or a membrane hairpin topology, containing two TMDs (Ruch and Machamer, 2012).

Both SARS-CoV E and SARS-CoV-2 E shared a 91% sequence identity with the amino acid sequence of template 5×29, suggesting high sequence homology (Figures S1A and S1B). This is to be expected given the 94% sequence similarity between SARS-CoV E and SARS-CoV-2 E, and since 5×29 spans residues 8-65 of the SARS-CoV E UniProt reference sequence (P59637), which excludes the PBM (Grifoni, et al., 2020; Schoeman and Fielding, 2020). The RMSD analysis showed a 0.698Å and 2.155Å difference between SARS-CoV E and SARS-CoV-2 E and the template 5×29, respectively, suggesting very little structural deviation between the respective two structures and the template. The RMSD analysis between MERS-CoV E and the homologous template structure 2mm4 indicated 1.458Å difference also suggesting a slight amount of structural deviation between the two structures. The 3D structures predicted for the E proteins of SARS-CoV, SARS-CoV-2, and MERS-CoV, therefore, successfully satisfied the Ramachandran plot dihedral angle distributions and, based on the low RMSD values, the correct protein folds were assigned to the respective protein sequences.

Despite the moderate sequence identity between the target sequence of the hCoV-229E E protein and the 5×29 template, the protein model from this pairwise sequence alignment will provide a medium accuracy protein model which is still useful for domain comparisons. The RMSD analysis between hCoV-229E E and the homologous template structure 5×29 indicated a 1.776Å difference, suggesting very little deviation between the two structures. The RMSD analysis between hCoV-NL63 E and the homologous template structure (229E protein model) similarly indicated a 1.217Å difference, also suggesting very little deviation between the generated model and the homologous template structure. Thus, the 3D structures predicted for the E proteins of hCoVs 229E and NL63 successfully passed quality assessment and based on the low RSMD values, the correct protein folds were assigned to the respective protein sequences.

The RMSD analysis between hCoV-OC43 E and the homologous template structure 5×29 indicated 3.328Å difference, whereas the RMSD analysis between hCoV-HKU1 E and the homologous template structure 229E indicated a 5.343Å difference, both of which suggest a large deviation between the generated models and their respective homologous template structures. Although the 3D structure predicted for the hCoV-HKU1 E protein satisfied the Ramachandran quality check, a high RSMD value suggests the incorrect fold was assigned to the protein sequence.

The E protein structures of the less virulent hCoVs OC43 and HKU1 predicted by MODELLER had the lowest sequence identities and the highest RMSD values compared to their templates. The high RMSD values are due to misorientations of the C-terminal domain regions. This is a common limitation of homology modelling if sequence identity and coverage are low in a specific region resulting in low accuracy protein models. However, the ITASSER predicted protein models for OC43 and HKU1 displayed lower RMSD values when aligned to template 2mm4, although the PBM was not conserved and displayed a flexible coil region (Figures S3A and S3B). We are confident in the E protein models predicted using MODELLER that display a less flexible PBM for the less virulent hCoVs.

The flexibility of the region in which the E protein PBM is located appears to distinguish the more virulent hCoVs quite clearly from the less virulent hCoVs; the former containing a PBM with a more flexible coil region, while the latter is characterised by a PBM that contains a less flexible α-helical turn. While so far only five interaction partners have been identified for SARS-CoV E, more may yet be identified and those that have already been identified have very different structures (Jimenez-Guardeño, et al., 2014; Nieto-Torres, et al., 2011; Teoh, et al., 2010; Yang, et al., 2005). This could quite possibly be a pathogenic feature that allows the more virulent hCoVs to interact with a wider range of host proteins which could explain their increased pathogenicity and severity of the disease they cause. By comparison, the less flexible α-helical turn of the less virulent hCoV E protein PBM could very likely be an impeding characteristic, preventing or limiting interaction with host proteins, which could possibly explain the limited pathogenic capabilities of the less virulent hCoVs. Without experimental data, however, this remains to be determined.

Clearly, much is still unknown about the topology of the hCoV E protein and how different hCoVs can exhibit different topologies while still performing the same functions. However, this is the first study, to predict and generate 3D models of the E protein for all seven hCoVs. This study also attempted to determine a distinguishing feature between the E proteins of the more virulent and the less virulent hCoVs to provide a possible explanation for the difference in pathogenicity and disease severity. SARS-CoV is the only hCoV for which E protein interaction partners have been identified. It, therefore, remains to be determined whether the E protein of any of the less virulent hCoVs can also interact with host cell proteins similar to SARS-CoV E, and, if so, which host cell proteins the less virulent hCoV’s E proteins can interact with. As it stands, there is no experimental evidence on the topology of the E protein for any of the less virulent hCoVs, leaving questions to be answered: can one protein adopt multiple topologies, each one for a different function? If the E protein of less virulent hCoVs exhibits a topology opposite to that of the more virulent hCoVs, can the less virulent hCoVs still perform similar pathogenic functions like ion-channel activity and viral-host protein-protein interactions with a PBM? Answers to such questions could further help to explain the difference in severity between the different hCoVs and could provide insight into membrane protein topology- and structure-function relationships as well.

## Acknowledgements

We apologize to any author whose work has been inadvertently omitted from this article.

## Supplementary data

**Table S2.**
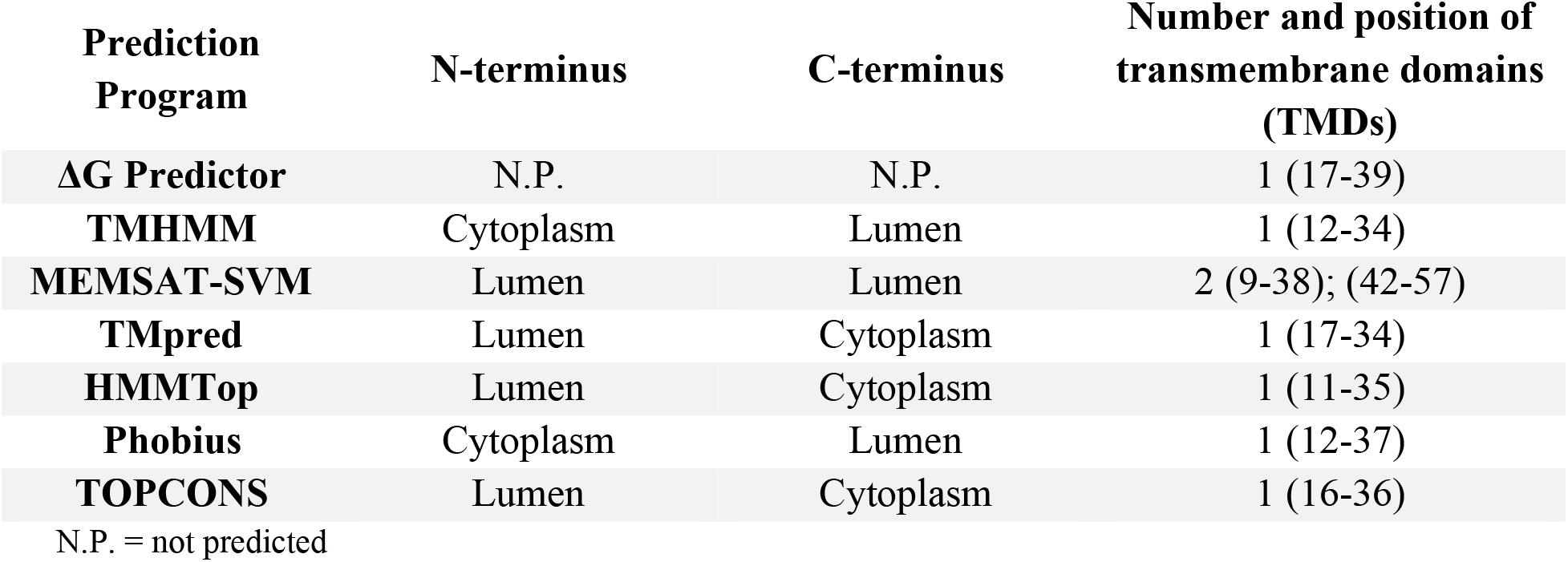
Predicted topologies of severe acute respiratory coronavirus (SARS-CoV) envelope (E) protein (Accession number: P59637) using prediction programs ΔG Predictor, TMHMM, MEMSAT-SVM, TMpred, HMMTop, Phobius, TOPCONS. For each program, the predicted location of the amino (N) and carboxy (C) terminus of the E protein is indicated as being in the lumen of the endoplasmic reticulum-Golgi intermediate compartment (ERGIC) or in the cytoplasm of infected host cells. The number of transmembrane domains (TMDs) and the corresponding length of residues that they span is also indicated as predicted by each program.

**Table S3.**
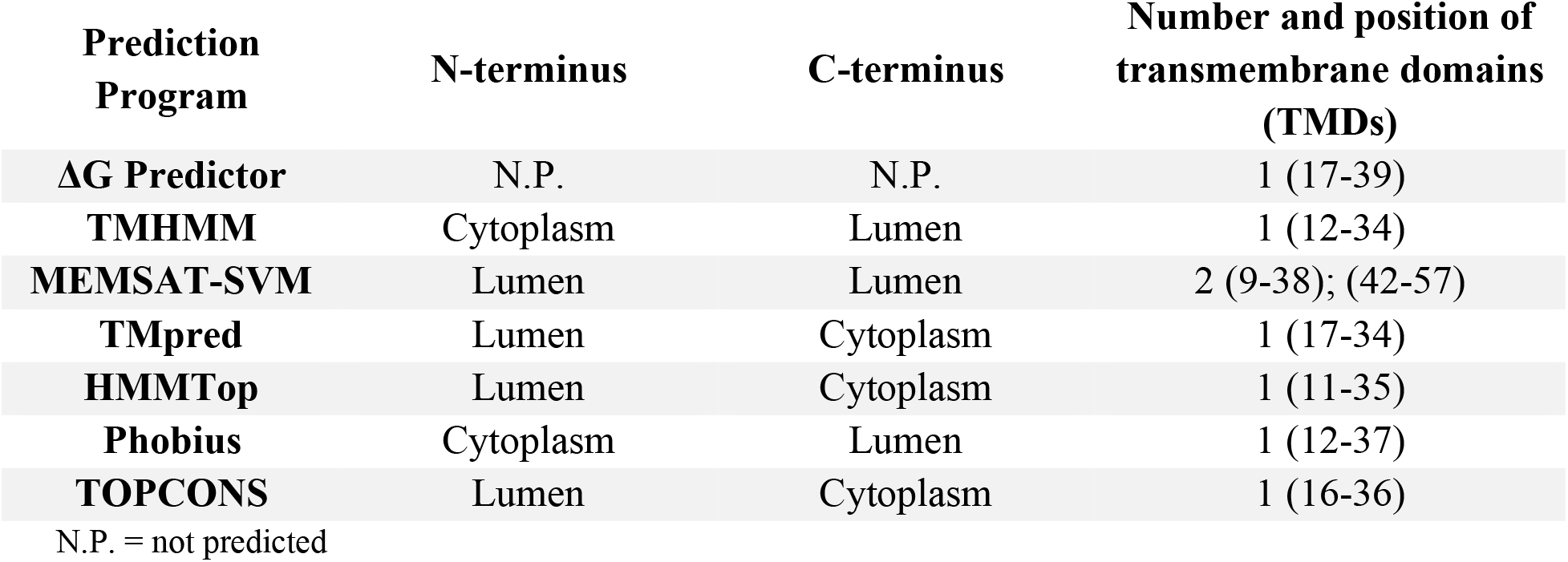
Predicted topologies of severe acute respiratory coronavirus 2 (SARS-CoV-2) envelope (E) protein (Accession number: P0DTC4) using prediction programs ΔG Predictor, TMHMM, MEMSAT-SVM, TMpred, HMMTop, Phobius, TOPCONS. For each program, the predicted location of the amino (N) and carboxy (C) terminus of the E protein is indicated as being in the lumen of the endoplasmic reticulum-Golgi intermediate compartment (ERGIC) or in the cytoplasm of infected host cells. The number of transmembrane domains (TMDs) and the corresponding length of residues that they span is also indicated as predicted by each program.

**Table S4.**
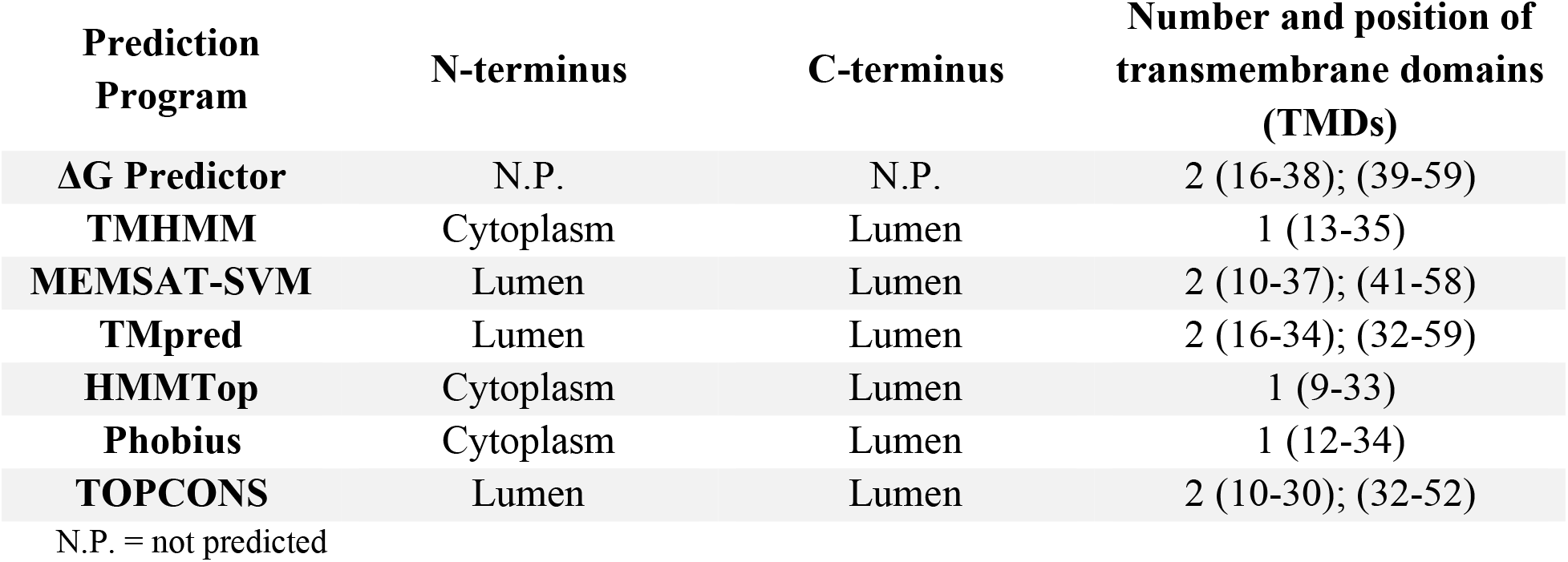
Predicted topologies of Middle East respiratory coronavirus (MERS-CoV) envelope (E) protein (Accession number: K9N5R3) using prediction programs ΔG Predictor, TMHMM, MEMSAT-SVM, TMpred, HMMTop, Phobius, TOPCONS. For each program, the predicted location of the amino (N) and carboxy (C) terminus of the E protein is indicated as being in the lumen of the endoplasmic reticulum-Golgi intermediate compartment (ERGIC) or in the cytoplasm of infected host cells. The number of transmembrane domains (TMDs) and the corresponding length of residues that they span is also indicated as predicted by each program.

**Table S5.** Predicted topologies of human coronavirus 229E (hCoV-229E) envelope (E) protein (Accession number: P19741) using prediction programs ΔG Predictor, TMHMM, MEMSAT-SVM, TMpred, HMMTop, Phobius, TOPCONS. For each program, the predicted location of the amino (N) and carboxy (C) terminus of the E protein is indicated as being in the lumen of the endoplasmic reticulum-Golgi intermediate compartment (ERGIC) or in the cytoplasm of infected host cells. The number of transmembrane domains (TMDs) and the corresponding length of residues that they span is also indicated as predicted by each program.

| Prediction Program | N-terminus | C-terminus | Number and position of transmembrane domains (TMDs) |
| --- | --- | --- | --- |
| **ΔG Predictor** | N.P. | N.P. | 1 (15-36) |
| **TMHMM** | Lumen | Cytoplasm | 1 (10-32) |
| **MEMSAT-SVM** | Cytoplasm | Lumen | 1 (12-42) |
| **Tmpred** | Lumen | Cytoplasm | 1 (15-33) |
| **HMMTop** | Cytoplasm | Lumen | 1 (11-33) |
| **Phobius** | Lumen | Cytoplasm | 1 (15-36) |
| **TOPCONS** | Lumen | Cytoplasm | 1 (13-33) |

N.P. = not predicted

**Table S6.** Predicted topologies of human coronavirus NL63 (hCoV-NL63) envelope (E) protein (Accession number: Q6Q1S0) using prediction programs ΔG Predictor, TMHMM, MEMSAT-SVM, TMpred, HMMTop, Phobius, TOPCONS. For each program, the predicted location of the amino (N) and carboxy (C) terminus of the E protein is indicated as being in the lumen of the endoplasmic reticulum-Golgi intermediate compartment (ERGIC) or in the cytoplasm of infected host cells. The number of transmembrane domains (TMDs) and the corresponding length of residues that they span is also indicated as predicted by each program.

| Prediction Program | N-terminus | C-terminus | Number and position of transmembrane domains (TMDs) |
| --- | --- | --- | --- |
| **ΔG Predictor** | N.P. | N.P. | 1 (16-36) |
| **TMHMM** | Lumen | Cytoplasm | 1 (10-32) |
| **MEMSAT-SVM** | Cytoplasm | Lumen | 1 (12-42) |
| **Tmpred** | Cytoplasm | Lumen | 1 (15-33) |
| **HMMTop** | Cytoplasm | Lumen | 1 (11-33) |
| **Phobius** | Cytoplasm | Lumen | 1 (12-33) |
| **TOPCONS** | Lumen | Cytoplasm | 1 (23-43) |

N.P. = not predicted

**Table S7.** Predicted topologies of human coronavirus OC43 (hCoV-OC43) envelope (E) protein (Accession number: Q04854) using prediction programs ΔG Predictor, TMHMM, MEMSAT-SVM, TMpred, HMMTop, Phobius, TOPCONS. For each program, the predicted location of the amino (N) and carboxy (C) terminus of the E protein is indicated as being in the lumen of the endoplasmic reticulum-Golgi intermediate compartment (ERGIC) or in the cytoplasm of infected host cells. The number of transmembrane domains (TMDs) and the corresponding length of residues that they span is also indicated as predicted by each program.

| Prediction Program | N-terminus | C-terminus | Number and position of transmembrane domains (TMDs) |
| --- | --- | --- | --- |
| **ΔG Predictor** | N.P. | N.P. | 2 (18-39); (41-61) |
| **TMHMM** | Lumen | Cytoplasm | 1 (15-37) |
| **MEMSAT-SVM** | Lumen | Lumen | 2 (12-39); (43-58) |
| **Tmpred** | Cytoplasm | Cytoplasm | 2 (15-33); (41-61) |
| **HMMTop** | Lumen | Lumen | 2 (18-36); (43-61) |
| **Phobius** | Lumen | Cytoplasm | 1 (12-39) |
| **TOPCONS** | Lumen | Lumen | 2 (16-36); (38-58) |

N.P. = not predicted

**Table S8.** Predicted topologies of human coronavirus HKU1 (hCoV-HKU1) envelope (E) protein (Accession number: Q5MQC8) using prediction programs ΔG Predictor, TMHMM, MEMSAT-SVM, TMpred, HMMTop, Phobius, TOPCONS. For each program, the predicted location of the amino (N) and carboxy (C) terminus of the E protein is indicated as being in the lumen of the endoplasmic reticulum-Golgi intermediate compartment (ERGIC) or in the cytoplasm of infected host cells. The number of transmembrane domains (TMDs) and the corresponding length of residues that they span is also indicated as predicted by each program.

| Prediction Program | N-terminus | C-terminus | Number and position of transmembrane domains (TMDs) |
| --- | --- | --- | --- |
| **ΔG Predictor** | N.P. | N.P. | 2 (17-39); (40-61) |
| **TMHMM** | Lumen | Lumen | 2 (10-32); (39-58) |
| **MEMSAT-SVM** | Lumen | Lumen | 2 (10-36); (40-55) |
| **TMpred** | Lumen | Lumen | 2 (16-34); (39-59) |
| **HMMTop** | Lumen | Lumen | 2 (17-36); (43-59) |
| **Phobius** | Lumen | Lumen | 2 (12-34); (39-59) |
| **TOPCONS** | Lumen | Lumen | 2 (10-30); (32-52) |

N.P. = not predicted

**Figure S3.**
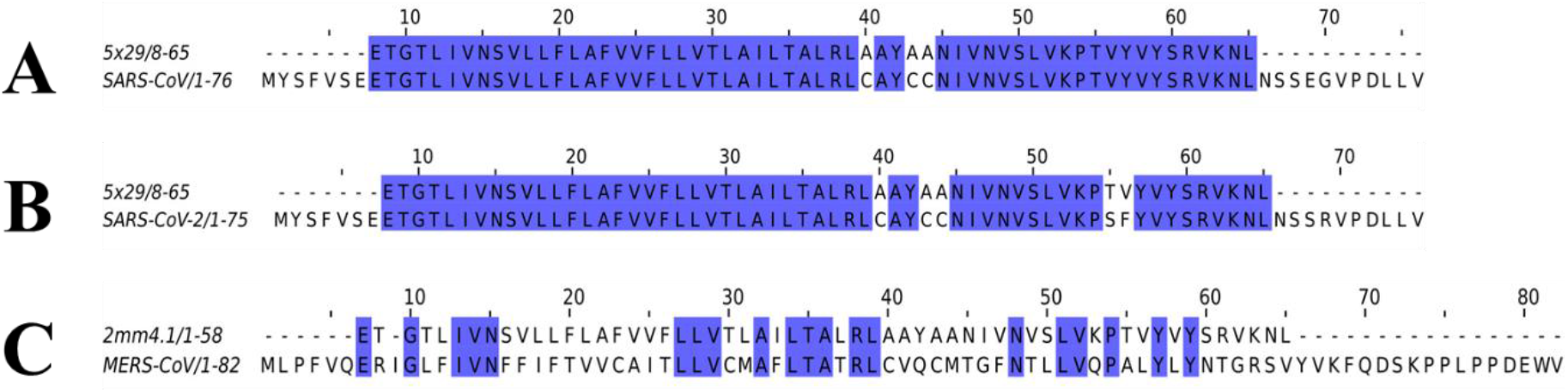
Pairwise sequence alignments between the more virulent human (h) coronavirus (CoV) envelope (E) proteins and the respective template used to generate the three-dimensional (3D) model. Sequence alignments were generated using Jalview (v2.11.1.3) and coloured by sequence identity (blue). **A.** Pairwise sequence alignment between the SARS-CoV E protein (P59637) and template 5×29. Sequences shared 91% identity with no conserved residues in the PDZ-binding motif (DLLV). **B.** Pairwise sequence alignment between the SARS-CoV-2 E protein (P0DTC4) and template 5×29. Sequences shared 91% identity with no conserved residues in the PDZ-binding motif (DLLV). **C.** Pairwise sequence alignment between the MERS-CoV E protein (K9N5R3) and template 2mm4. Sequences shared 35% identity with no conserved residues in the PDZ-binding motif (DEWV).

**Figure S4.**
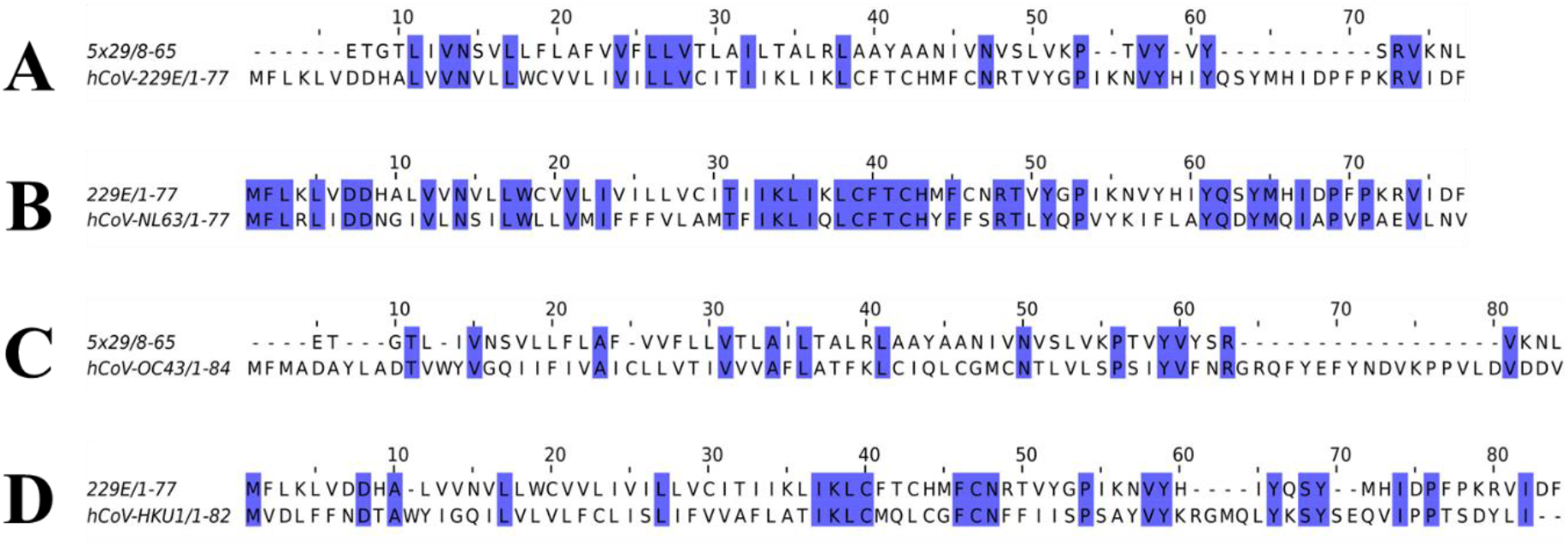
Pairwise sequence alignments between the less virulent human (h) coronaviruses (CoVs) and the respective template used to generate the three-dimensional (3D) model. Sequence alignments were generated using Jalview (v2.11.1.3) and coloured by sequence identity (blue). **A.** Pairwise sequence alignment between the hCoV-229E E protein (P19741) and template 5×29. Sequences shared 29% and PDZ-binding motif residues <u>V</u>IDF were conserved. **B.** Pairwise sequence alignment between the hCoV-NL63 E protein (Q6Q1S0) and the hCoV-229E E protein homologous structure. Sequences shared 47% identity and PDZ-binding motif residues <u>V</u>LNV were conserved. **C.** Pairwise sequence alignment between the hCoV-OC43 E protein (Q04854) and template 5×29. Sequences shared 22% identity and PDZ-binding motif residues <u>V</u>DDV were conserved. **D.** Pairwise sequence alignment between the hCoV-HKU1 E protein (Q5MQC8) and the hCoV-229E E protein homologous structure. Sequences shared 27% identity and PDZ-binding motif residues DYL<u>I</u> were conserved.

**Figure S5.**
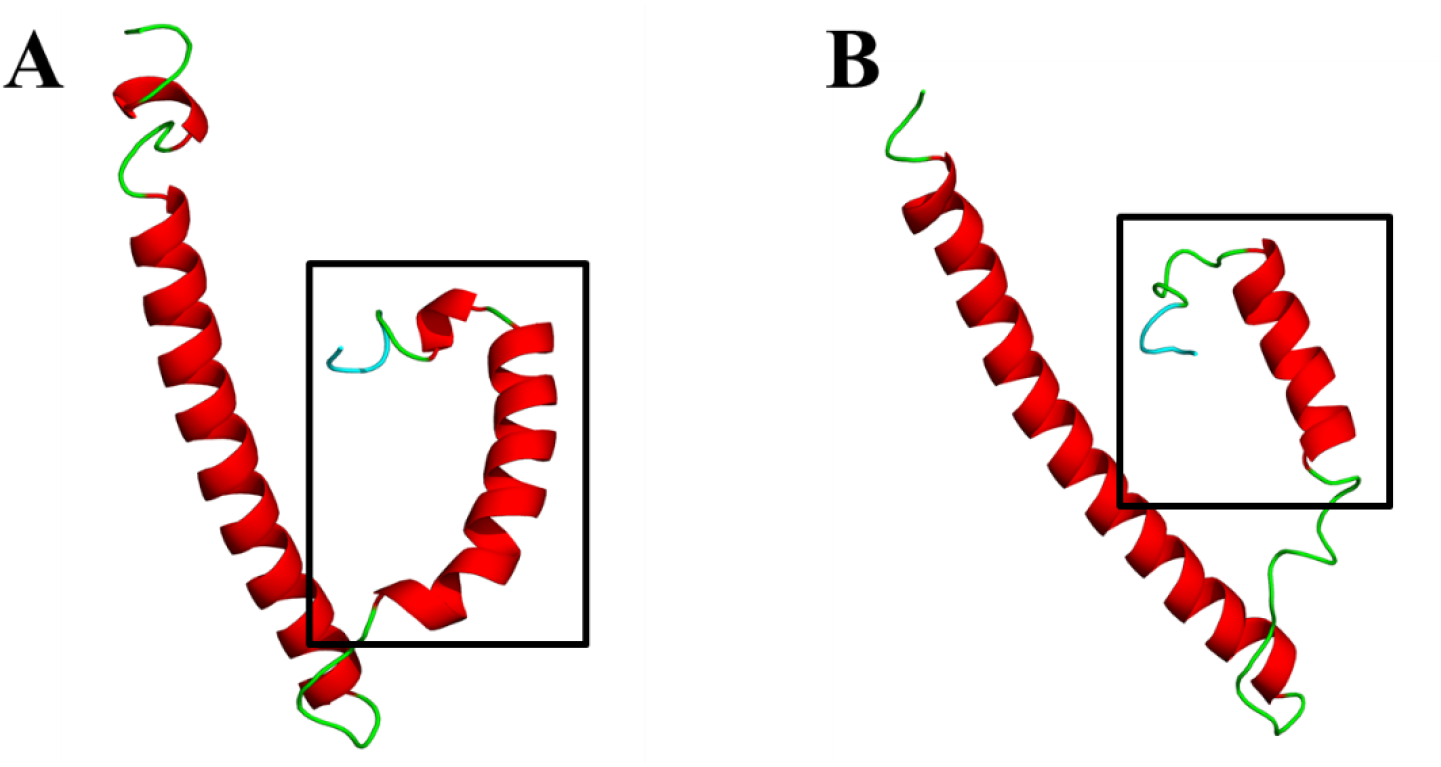
Cartoon representation of the three-dimensional (3D) models of the envelope (E) protein of hCoV-OC43 and hCoV-HKU1. **A.** hCoV-OC43 E. **B.** hCoV-HKU1 E. Models were generated using ITASSER Webserver and based on the nuclear magnetic resonance (NMR)-resolved structure for SARS-CoV E (PDBID: 2mm4) obtained from the protein data bank (PDB) (Li et al., 2014). The PDZ-binding motif (PBM) is coloured in cyan and the squared region shows the variable C-terminal domain.

